# Impact of anaesthesia on static and dynamic functional connectivity in mice

**DOI:** 10.1101/2021.03.18.436098

**Authors:** Tomokazu Tsurugizawa, Daisuke Yoshimaru

## Abstract

A few studies have compared the static functional connectivity between awake and anaesthetized states in rodents by resting-state fMRI. However, impact of anaesthesia on static and dynamic fluctuations in functional connectivity has not been fully understood. Here, we developed a resting-state fMRI protocol to perform awake and anaesthetized functional MRI in the same mice. Static functional connectivity showed a widespread decrease under anaesthesia, such as when under isoflurane or a mixture of isoflurane and medetomidine. Several interhemispheric connections were key connections for anaesthetized condition from awake. Dynamic functional connectivity demonstrates the shift from frequent broad connections across the cortex, the hypothalamus, and the auditory-visual cortex to frequent local connections within the cortex only. Fractional amplitude of low frequency fluctuation in the thalamic nuclei decreased under both anaesthesia. These results indicate that typical anaesthetics for functional MRI alters the spatiotemporal profile of the dynamic brain network in subcortical regions, including the thalamic nuclei and limbic system.

**Highlights:**

- Resting-state fMRI was compared between awake and anaesthetized in the same mice.
- Anaesthesia induced a widespread decrease of static functional connectivity.
- Anaesthesia strengthened local connections within the cortex.
- fALFF in the thalamus was decreased by anaesthesia.

## Introduction

The effect of anaesthetics on the brain functional network has interested researchers for decades. Functional connectivity (FC), which reflects the synchronization of fluctuation of blood oxygenation level-dependent (BOLD) signals accompanying neuronal activity among anatomically separated brain regions ^1^, could be a key structure between awake and anaesthetized states ^2^. A common assumption is that FC does not change over the data acquisition period, which is generally between 5 and 20 min, and therefore, static FC has been computed from the time course of BOLD signal throughout the acquisition time. Previous studies have shown that static FC is widely decreased by anaesthesia in humans ^3, 4^ and in monkeys ^5^, depending on the agent, dose, and target functional network. Rodent models are promising for deeply understanding the effect of anaesthesia on FC, and recent progress in fMRI has enabled the investigation of resting-state functional connectivity in awake mouse ^6, 7, 8^, as well as rats ^9, 10^. Awake mouse fMRI data indicates that connectivity strength is significantly decreased in widespread brain regions under anaesthesia ^11, 12, 13^, but the structure of functional connectivity between awake and anaesthetized animals is similar ^14^.

Although static FC studies reflect the synchronization of neuronal activity, FC is not absolutely stationary in nature during acquisition. This concept of dynamic fluctuation of functional connectivity, which is called dynamic functional connectivity (dynamic FC), has emerged in the last 10 years ^15^. Dynamic FC is altered in psychiatric disease patients ^16, 17, 18^, and recent studies have shown the modulation of dynamic FC by anaesthetics in macaque models ^5^. The dominating functional configurations have low information capacity and lack negative correlations under anaesthesia, and conversely, dynamical exploration of a rich, flexible repertoire of functional configurations in awake state ^19^. A rodent study has also shown that isoflurane (Iso)-anaesthesia profoundly impacted the dynamic FC of neural circuits subserving cognitive and emotional functions by investigating the FC correlated to the seed regions of the infralimbic cortex and somatosensory cortex ^20^. However, the effect of anaesthesia on the dynamic properties of other functional networks, such as default mode network (DMN) and cortical network, is not known.

In the current study, we aimed to investigate the differences in static and dynamic functional connectivity under anaesthesia and in the awake state. For this purpose, we successfully induced anaesthesia in awake mice in a magnetic bore. This enabled us to compare the awake state and anaesthetized state in the same group directly. Two commonly used anaesthesia protocols for fMRI, a low dose of Iso and a mixture of low doses of Iso and medetomidine (Med), were used in this study. Furthermore, fractional amplitude of low frequency fluctuation (fALFF) was investigated to assess the alteration of the amplitude of neuronal activity at the voxel level ^21^.

## Method

### Animals and surgery

We used 27 male C576J/BL mice (18 - 25 g), allocated to 9 Iso group, 9 Iso + Med group, and 9 test-retest reproducibility group. All animal procedures used in the present study were approved by the *Ethical Committee for Animal Experiments* (*Comité d’Ethique en Expérimentation Animale, Commissariat à l’Energie Atomique et aux Énergies Alternatives, Direction des Sciences du Vivant*, Fontenay-aux-Roses, 3 France). All methods in this study were performed in accordance with the relevant guidelines and regulations.

### Surgery for awake fMRI

We performed cranioplasty on the mice under isoflurane anaesthesia (1.5% with air) for awake fMRI ^22^. The skin of the head of the mice was cut longitudinally, and the skull surface was exposed and polished using saline and gauze. We applied resin cement (Super-Bond C&B, Sun Medical Company, Ltd., Shiga, Japan) on the surface of the skull. Then, a cranioplastic acrylic head bar (3 × 3 × 27 mm^3^), which was used for head fixation without pain during the imaging session, was attached to the skull horizontally with resin cement (Unifast TRAD, GC Corporation, Tokyo, Japan). Following surgery, the mice were allowed to recover for more than 1 week (Fig. 1). (Fig. 1)

**Figure 1.**
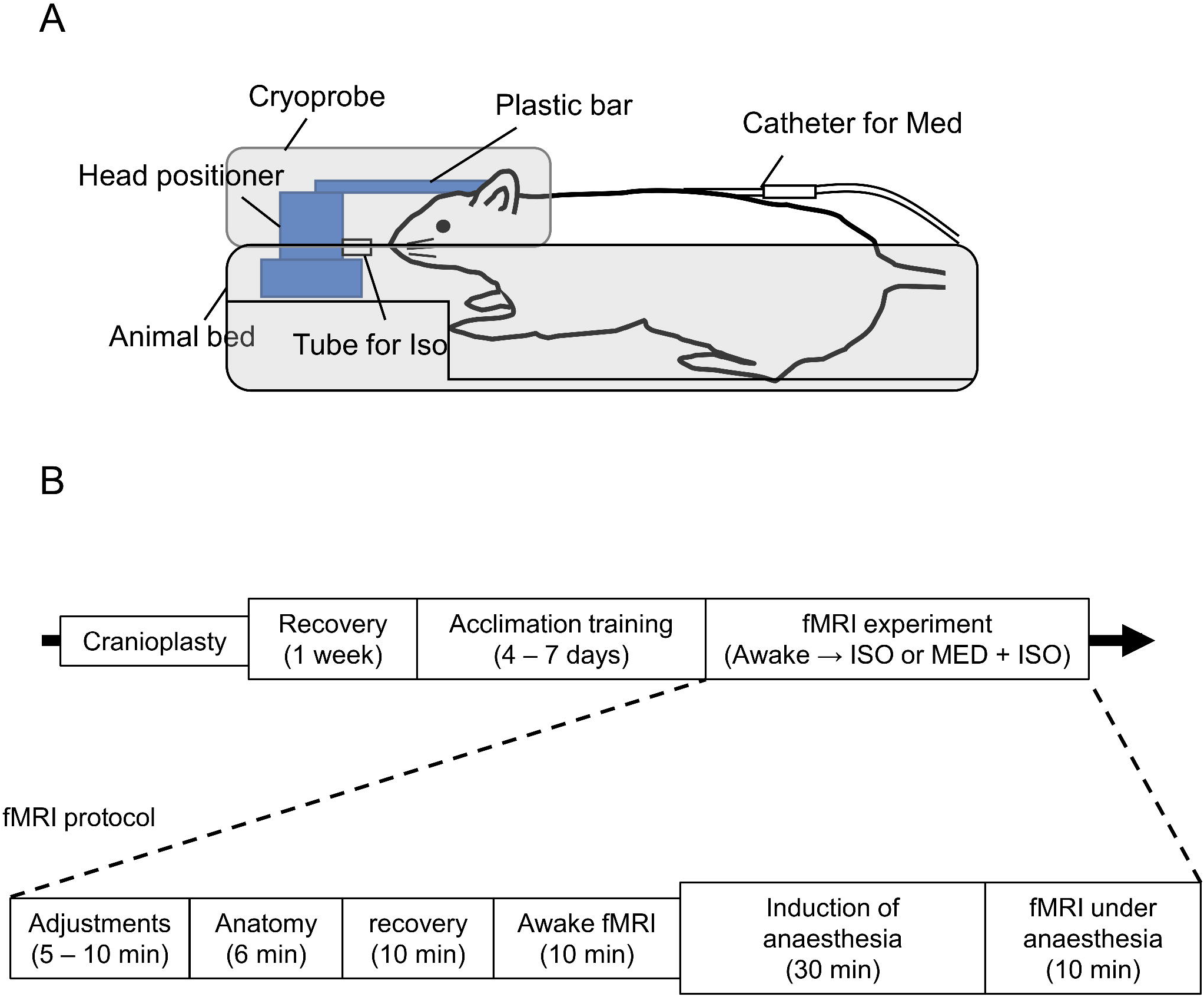
Schematic figure of the experimental paradigm. (A) Tube for Iso-anaesthesia is placed in front of the nose and catheter for Med injection was implanted under the back skin. The plastic bar was attached on the skull by dental cement was fixed by head positioner. (B) Schedule of experiment.

### fMRI acclimation training

During surgery recovery, mice were allowed to become accustomed to the cupped handling method ^7, 23^. We confirmed that mice did not urinate and did not jump from the cupped hands during handling before acclimation training. The mice were trained for 4-6 days to become accustomed to the fMRI conditions. During the first 2 days, a pseudo-MRI system was used. The mice were anaesthetized by 1.5% isoflurane for maintenance while positioning them on the pseudo-MRI apparatus. Mice remained in the pseudo-MRI apparatus for 30 min on the first day and 90 min on the second day. The respiratory rate of the mice was monitored during the training sessions. For the next 2-4 days, mice were placed on an MRI bed with a head fixation system with the same methods as the fMRI experiment (see the awake fMRI section for details about awake fMRI procedure). Throughout the training period, the respiration rate and body weight were confirmed to be in the normal range using an MR-compatible monitoring system (model 1025; SA Instruments, Stony Brook, NY, USA).

### Awake and anaesthetized fMRI

All MRI experiments were performed with a Bruker 11.7 T MRI system with a cryoprobe (Bruker BioSpin, Ettlingen, Germany). The homemade head fixation system was placed on the mouse bed for cryoprobe ^22^. The mice were anaesthetized by 2% isoflurane with air for induction and less than 1.5% with air for maintenance during head fixation and adjustment of the magnetic homogeneity, shimming and reference pulse. The schematic figure of anaesthesia system is shown in Fig. 1A. Iso was administered for no more than 10 min. The anaesthesia was stopped and structural image scan started. The mice awoke during 10 min recovery period. We confirmed that the respiration rate increased over 150 breaths/min during the recovery period. Previous study showed that neuronal activity recovered to that of a conscious state at 10 min after cessation of Iso delivery ^7^. The fMRI acquisition started at that point. The schematic paradigm of the MRI protocol is illustrated in Fig. 1B. When mice awakened from anaesthesia after the recovery period, awake resting-state fMRI was acquired. Once awake fMRI was complete, anaesthesia was induced. In the Iso group, 1.5% isoflurane with air was delivered for the induction of anaesthesia through the tube (Fig. 1A). When respiration was decreased to less than 100/min, isoflurane was reduced to 0.8%. fMRI scanning started 30 min after isoflurane induction was initiated. For the Iso + Med group, 1.5% isoflurane with air was delivered for the induction of anaesthesia. When the respiration rate decreased to less than 100/min, a bolus of medetomidine (0.1 mg/kg body weight, s.c.) was injected through the infusion canula, which was inserted under the skin on the animals’ back, and then it was infused continuously at a rate of 0.05 mg/kg body weight/h (s.c.). Isoflurane was reduced to 0.5%when the respiratory rate was reduced to less than 100/min. The resting state fMRI images were acquired using a T2*-weighted multi-slice gradient-echo EPI sequence with the following parameters: TR/TE = 1500/14 ms, spatial resolution = 100 x 100 x 500 µm^3^/voxel, 15 slices and 400 volumes (total 10 min). For normalization in image processing, the structural image with the same field of view as the fMRI was acquired using T2-weighted multi-slice rapid acquisition with relaxation enhancement (RARE) with the following parameters: TR = 2,500 ms, effective TE = 13 ms, spatial resolution = 100 x 100 x 500 µm^3^, RARE factor = 4, and 4 averages.

### Image preprocessing

Statistical parametric mapping SPM12 software (Wellcome Trust Center for Neuroimaging, UK) was used to analyse the fMRI data and to perform preprocessing steps, including slice timing correction, motion correction by realignment, normalization and smoothing with a Gaussian filter (0.3 x 0.3 mm^2^ in each slice). Before preprocessing, template images co-registered to the Allen mouse brain atlas were obtained (http://atlas.brain-map.org/). Functional and structural images were normalized to these template images. Framewise displacement (FD) was used to check head motion ^24^. FD was calculated using 6 motion parameters (3 for translation and 3 for rotation) from realignment in SPM. For all subjects, it was confirmed that the head motion was less than the following criteria: (1) mean FD averaged for all time points during the scan was less than 0.02 mm and (2) FDs at all time points were less than 0.05 mm. Mean FD was not significantly different between groups. The linear trend in the time course of the fMRI dataset was removed voxel-by-voxel using the CONN toolbox (https://web.conn-toolbox.org/) ^25^. Mean signals in the ventricles and white matter and six motion parameters of an object (translational and rotational motions) were regressed out from the time series of each voxel to reduce the contribution of physiological noise, such as respiration and head movement.

### Static functional connectivity

The preprocessed fMRI data were then detrended, and slow periodic fluctuations were extracted using a bandpass filter (0.01–0.08 Hz). A total of 142 regions of interest (ROIs, 71 per hemisphere) were delineated based on the template images, with reference to the Allen mouse brain atlas (https://scalablebrainatlas.incf.org/mouse/ABA12). The correlation coefficients between two ROIs were calculated using the CONN toolbox. The statistical significance of ROI-ROI connectivity was assessed by paired t-test or two sample t-test with a threshold of p < 0.05 (NBS-corrected), using the Network-Based 5 Statistic toolbox (https://www.nitrc.org/projects/nbs/). The connections with a Cohen’s d value exceeding 0.7 (minimum meaningful effect size) were further assessed for NBS. Subnetworks were then identified in the set of supra-threshold connections. The size of each subnetwork was measured in terms of the number of connections it comprised. Permutation testing was used to compute a family-wise error-corrected p-value for each subnetwork. Specifically, 5000 permutations were generated in which group labels were randomly permuted and the size of the largest subnetwork was recorded for each permutation. Subnetworks deemed to be significant with permutation testing (p < 0.05, family-wise error corrected) were identified as significant.

### Feature extraction

The feature extraction of alterations of functional connectivity was computed using MATLAB (MathWorks, Natick, USA). The least absolute shrinkage and selection operator (LASSO) method, which is suitable for the reduction of high-dimensional data, was used to select the significant features from the Pearson’s correlation coefficient data (10011 data sets) of regionally averaged fMRI time series obtained between all 142 brain regions. LASSO is a penalization method that shrinks all regression coefficients and sets the coefficients of many unrelated features that have no discriminatory power between the classes exactly to zero with the regulation parameter λ ^26, 27^. The regulation parameter λ tuned the sparsity of the model such that a larger λ would lead to a sparser model. We then selected the optimal λ in the LASSO model by using leave-one-out cross-validation via MSE plus one standard deviation of the MSE.

### The similarity of Functional Brain Connectivity

The differences among groups, individuals, and states of anaesthesia were directly examined by calculating the similarity among each original functional connectivity using MATLAB ^28, 29^. The similarity matrix was created by calculating Fisher’s z-transformed correlation between the linearized original connectivity matrices of each individual. This similarity matrix represents the awake group (n = 9) in the first third, the Iso group (n = 9) in the next third, and the Iso + Med group (n = 9) in the last third.

### Independent component analysis

Preprocessed fMRI data were analysed in the SPM Group ICA of fMRI Toolbox (GIFT v3.0b http://trendscentre.org/software/gift/). First, all data sets underwent a subject-specific principal component analysis (PCA), which estimated 150 components. All subjects’ reduced data sets were then concatenated and underwent a PCA that estimated 40 components at the group level. Subsequently, group-level spatial ICA was performed on the PCA output, identifying 40 functional components. The Infomax algorithm was used for component estimation. Subject-specific component maps and time courses were obtained from the group-level components using the GICA back reconstruction method. Using component spatial maps that we computed before ^7^, 27 components were finally identified as functional components. These components were the default mode network (DMN), including the retrosplenial cortex and the cingulate cortex (3 ICs); lateral cortical network (LCN), including the sensorimotor cortex (8 ICs); subcortical basal ganglion network (BG), including the striatum and the nucleus accumbens (2 ICs); hippocampus (Hip) (3 ICs); thalamic network (2 ICs); hypothalamus (Hypo) (4 ICs); and auditory-visual network (AUD-VIS), including the auditory and visual cortex (5 ICs).

### Dynamic FC

Dynamic FC was conducted with the GIFT Dynamic FNC Toolbox (v1.0a). We used time windows of 45 s (30 TRs) and steps of 1.5 s (1 TR). The time series was convolved with a Gaussian of sigma = 3 TRs. For each window, FC was estimated in the form of a regularized inverse covariance matrix using the Toolbox graphical LASSO method with an additional L1 norm constraint. The estimated covariances were transformed to Fisher-Z. To identify FC states that reoccurred across time and across subjects, the windowed FC matrices were subjected to the GIFT k-means clustering procedure. In the first step, a subset of windows (i.e., “exemplars”) was selected for each subject, representing those FC matrices with maximal variability in FC. From those windows, the optimal number of clusters (k) was determined by the toolbox using the elbow criterion, defined as the ratio of within-cluster distances to between-cluster distances. The resulting k cluster centroids were used as templates for clustering all windows’ FC matrices of all subjects. The dwell time of state was measured as the frequency of unchanged between time window and time+1 window, and the frequency of occurrence was measured as the number of windows in each state.

### Fractional amplitude of low frequency fluctuation

The fALFF was defined as the ratio of power within the frequency range between 0.01 and 0.08 Hz and power over the total frequency range in each voxel ^21, 30^. The fALFF maps were then compared, and significant differences between groups were tested voxel-by-voxel using paired t-tests or two-sample t-tests with standard progressive matrices with a threshold of p < 0.05 (false discovery rate (FDR) at the cluster level).

## Results

### Widespread reduction of static FC under each anaesthesia condition compared to the FC in the awake state

First, we investigated static FCs in awake, Iso anaesthetized and Iso + Med anaesthetized conditions. To compare awake and anaesthetized state directly, we developed the protocol to induce anaesthesia in awake mouse within MR bore (Fig. 1). Iso-anaesthesia was induced via the tube placed in front of the nose and Med was injected through the catheter under the back skin (Fig. 1A). Because the head space among the head fixation system, the mouse bed, and the cryoprobe is small enough to accumulate Iso, mice were anaesthetized in a few minutes following the start of Iso anaesthesia. The head fixation system described in the previous article was used in this study ^22^. We confirmed that mouse was anaesthetized when the respiratory rate was decreased from 150-200 beat/min ^7^, to less than 100 beat/min. To provide a whole-brain characterization of static FC, the whole brain was parcellated into 142 broad regions, and the Pearson correlation coefficient was calculated between pairs of regionally averaged fMRI time series to yield a 142 × 142 connectivity matrix in each mouse. The static FCs under Iso anaesthesia and Iso + Med anaesthesia conditions were widely decreased compared with those in the awake state (Fig. 2A). We noted that the typical functional structure of high interhemispheric connectivity (e.g., connectivity between the left sensorimotor cortex and right sensorimotor cortex) and the local network (e.g., within the sensorimotor cortex) was observed under Iso and Iso + Med anaesthesia conditions as well as in the awake state. To evaluate the reproducibility of awake fMRI, test-retest reproducibility was assessed. Awake fMRI was performed on two separate days, and static FCs were compared between the first and second days. There was no significant difference in static FC between 1st day and 2nd day (Supplementary Figs. 1A and 1B). These results and those of our previous study of awake fMRI ^7^ indicate excellent test-retest reproducibility. The network-based statistic (NBS) ^31^ was used to detect significant differences in functional connectivity strength in the pairs of brain regions between the awake and Iso-anaesthetized states and between the awake and Iso + Med anaesthetized states while controlling the familywise error rate. The statistical analysis shows widespread hypoconnectivity under Iso anaesthesia (Fig. 2B) and under Iso + Med anaesthesia (Fig. 2C). We did not observe a significant difference in static FC in pairs of regions between Iso and Iso + Med anaesthesia with an NBS-corrected significance level of p < 0.05 (Supplementary Fig. 1C). Independent component analysis (ICA) was used to investigate the typical network defined by previous studies in each state. We found that DMN, lateral cortical network (LCN), subcortical basal ganglion network (BG), hippocampus (Hip), thalamus (ThN), hypothalamic network (Hypo) and auditory-visual cortical network (AUD-VIS) were part of the network involved in awake, Iso anaesthetized and Iso + Med anaesthetized states (Supplementary Fig. 2).

**Figure 2.**
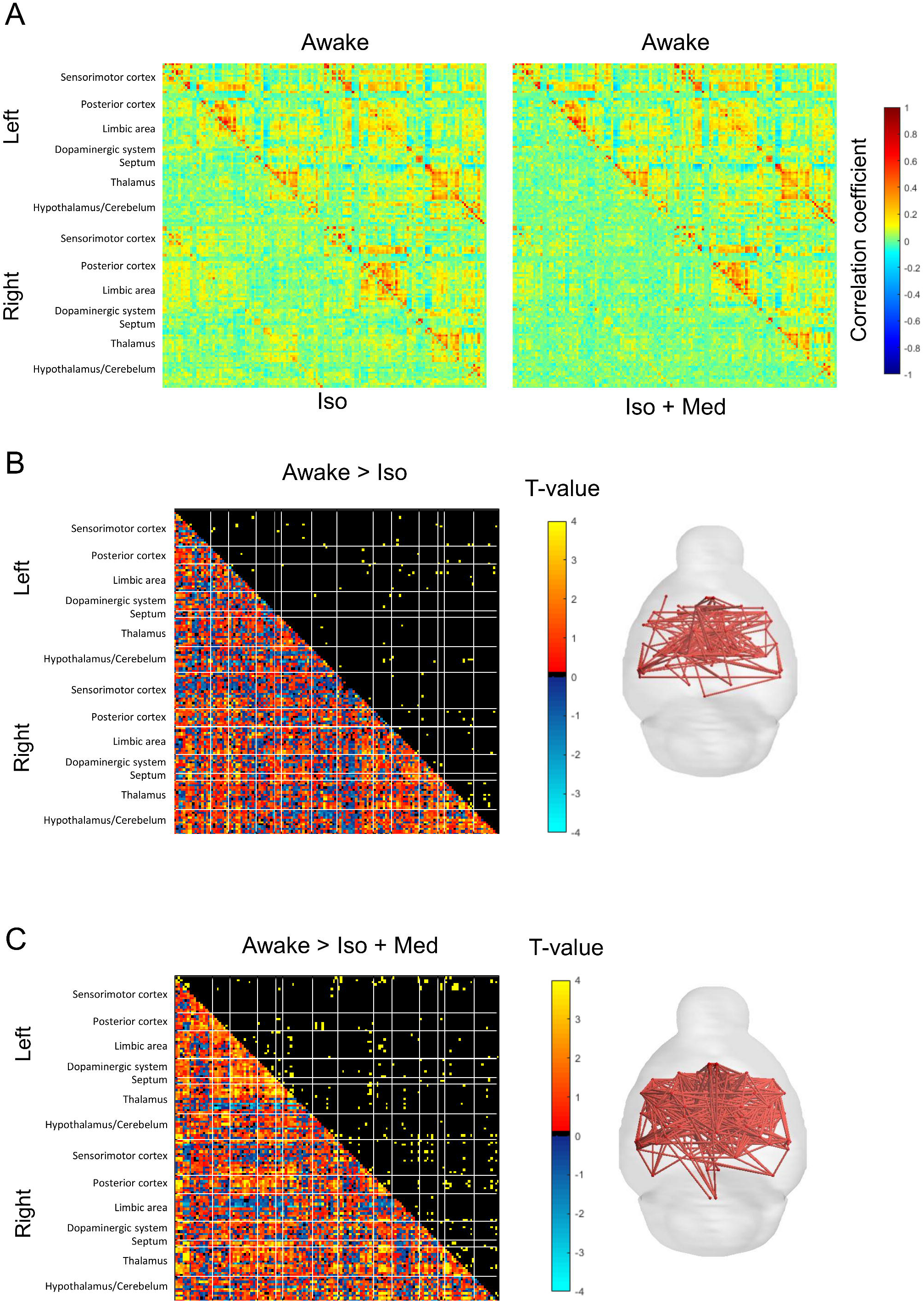
Comparison of static FC among awake, Iso-anaesthetized and Iso + Med-anaesthetized states. (A) Averaged connectivity matrices in awake, Iso anaesthetized, Iso + Med anaesthetized conditions. (B left) Comparison of each ROI-ROI correlation coefficient between awake and Iso groups. The lower left shows the T-values, and the upper right shows the significant differences (p < 0.05, NBS-corrected). (B right) Figure showing significantly decreased connectivity in the Iso-anaesthetized state compared with the awake state (p < 0.05, NBS-corrected). (C left) Comparison of each ROI-ROI correlation coefficient between the awake and Iso groups. The lower left shows the T-values, and the upper right shows the significant differences (p < 0.05, NBS-corrected). (C right) Figure showing significantly decreased connectivity in the Iso group compared with the Awake group.

We then investigated the similarity of static FC across all subjects in each condition (Fig. 3). High similarity was observed in the awake condition, whereas lower similarity was observed under both Iso and Iso + Med anaesthesia, indicating the robustness of the functional connectivity in the awake state and probably due to the reduction of the FC in the anaesthetized states. According to the similar static FC between Iso and Iso + Med anaesthesia, the anaesthetized state was defined as the combination of Iso and Iso + Med anaesthetized states for further static FC analysis.

**Figure 3.**
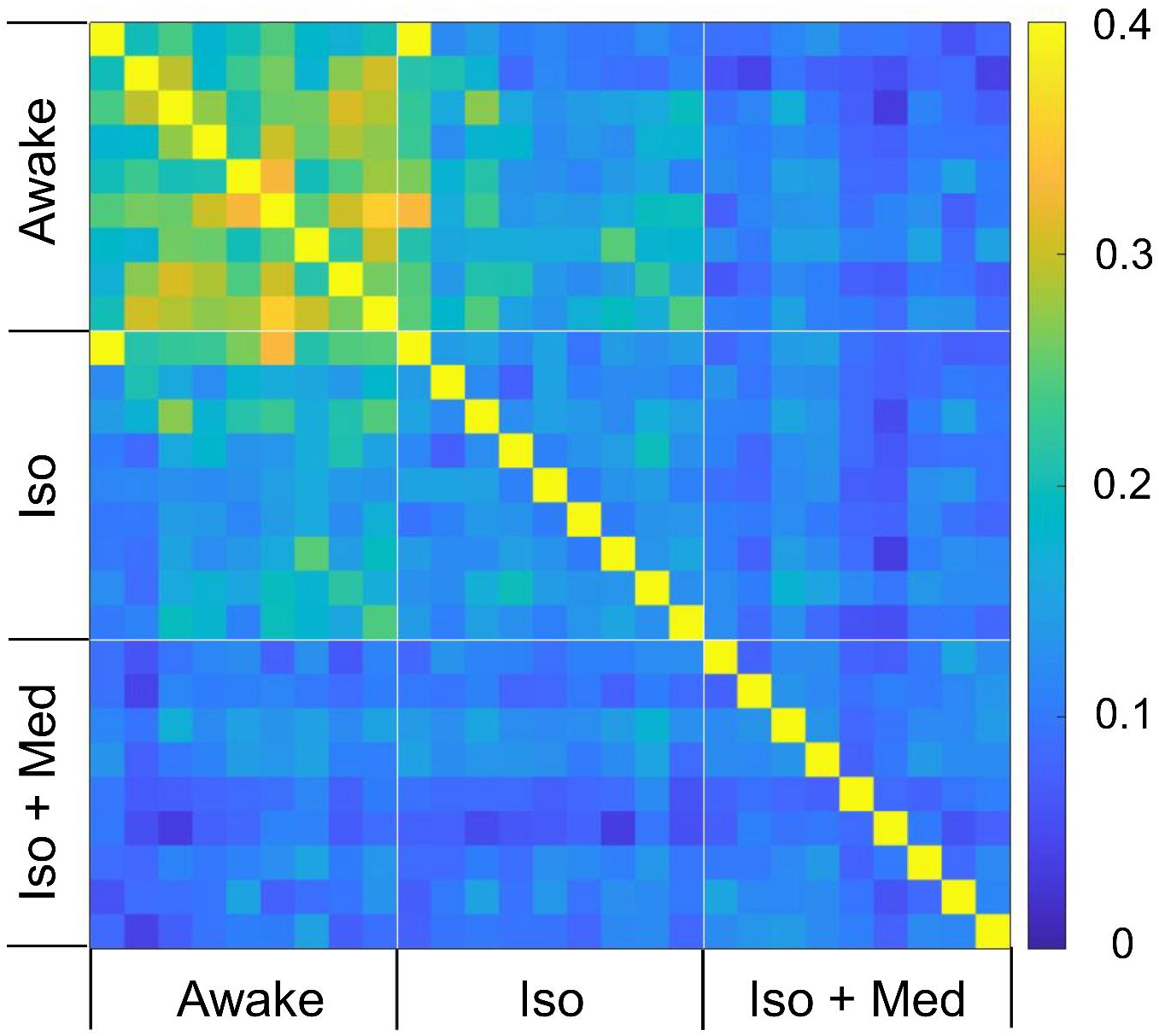
The similarity of functional brain connectivity among groups. A similarity matrix, where every cell represents the similarity of functional networks between different individuals. Rows and columns represent the similarity between a particular individual and all individuals, including other groups. The similarity matrix comprises the awake group (n = 9), Iso group (n = 9), and Iso + Med group (n = 9). Colour bar, correlation coefficients.

### Key nodes of widespread reduction of static FC under anaesthesia

Next, we tried to identify the key regions for widespread reduction of static FC under anaesthesia so that fMRI data under Iso and Iso + Med anaesthetized states were combined and evaluated as the anaesthesia group. Static FC under the anaesthetized state showed widespread hypoconnectivity compared to the FC in the awake state (Fig. 4A). The pairs of connections (a total of 142 x 141/2 = 10,011 connections) were then sequenced according to effect size (Cohen’s d), which was calculated from t-values (Fig. 4A). The top twenty connections (Cohen’s d > 1.0) demonstrated decreased static FC in the interhemispheric periaqueductal grey (PAG), anterior cingulate cortex (ACC), lateral septal nucleus (LSr), dorsomedial hypothalamus (DMH), (dorsal) posterior hypothalamus (PH, PHd), lateral hypothalamic area (LHA), secondary motor cortex (MOs), and dorsal agranular insular cortex (AId) (Fig. 4B). These results indicate that a significant reduction in interhemispheric connections in the whole brain could be a key connection under anaesthetization. Extensive reductions in connectivity strength according to effect size were observed in the whole brain, thalamic nuclei (ventral anterior-lateral complex (VAL), ventral posteromedial nucleus (VPM), ventral posterolateral nucleus (VPL), posterior complex (PO), centromedian nucleus (CM), ventral medial nucleus (VM)), limbic system (CA3 of hippocampal regions, LSr, lateral amygdala (LA) and central amygdala (CEA)), hypothalamus (LHA, medial preoptic area (MPO)), cortex (dorsal agranular insular cortex (AId), posterior agranular insular cortex (AIp), gustatory cortex (GU), secondary somatosensory cortex (SSs), dorsolateral entorhinal cortex (ENTDL), perirhinal cortex (PRh), piriform cortex (PIR), and anterior cingulate cortex (ACC)) (Fig. 4C). Then, we estimated the features of anaesthesia compared with those of the awake state. Based on the least absolute shrinkage and selection operator (LASSO) logistic regression model, the λ value that had the minimum mean squared error (MSE) was chosen as the optimal regulation parameter (λ = 0.1126), and 10 features with significant predictive power were selected using the regulation parameter from among all 10011 features (Figs. 4D and 4E).

**Figure 4.**
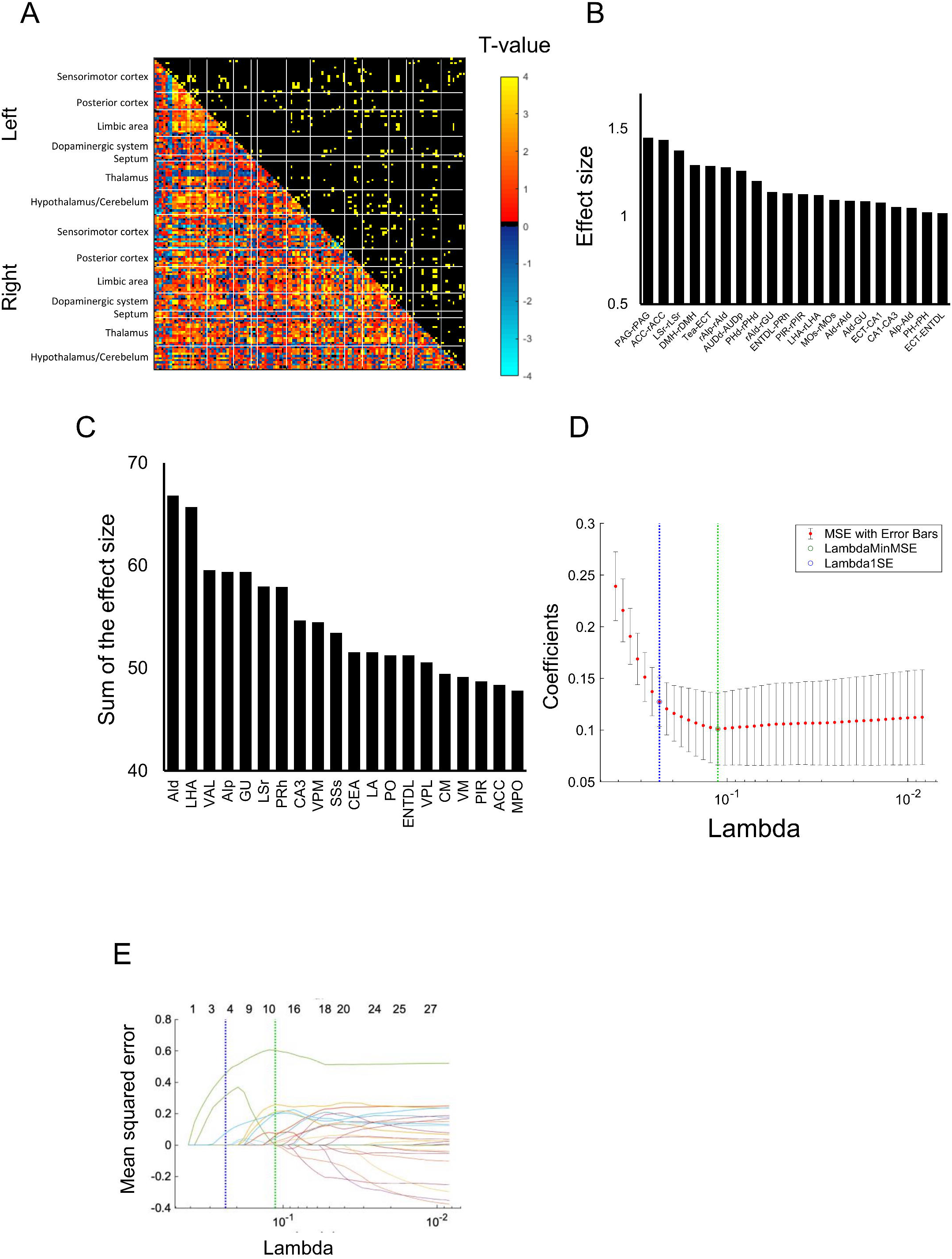
Widespread decrease in correlation coefficients under anaesthesia. (A) T-statistic values of each ROI-ROI correlation coefficient between awake and anaesthetized (combining Iso and Iso + Med anaesthesia) states. A positive t-value indicates higher correlation coefficients for the awake state than for the anaesthetized state. The upper right triangle shows the significant increase (yellow) or decrease (blue) (NBS-corrected). Colour bar, t-values. (B) Top twenty largest effect sizes of the ROI-ROI correlation coefficient. The effect sizes indicate a reduction in the correlation coefficient under the anaesthetized state. (C) Extent to which individual regions are affected by the network of reduced connectivity under anaesthetized conditions. The effect sizes were calculated across all connections associated with a given region. (D) A coefficient profile plot was created to visualize the cross-validated error of various levels of regularization. A vertical line was drawn at the value selected using leave-one-out cross-validation. The green circle and dotted line indicate the lambda with minimum cross-validation error. The blue circle and dotted line indicate the point with minimum cross-validation error plus one standard deviation. (E) The regulation parameter (λ) for the LASSO model was selected, and the trace plot of LASSO fit was visualized. Each line represents a trace of the values for a single predictor variable. PAG, periaqueductal grey; ACC, anterior cingulate cortex, LSr, lateral septal nucleus; DMH, dorsomedial hypothalamus; Tea, temporal association areas; ECT, ectorhinal area; AIp, posterior agranular insular cortex; AId, dorsal agranular insular cortex; AUDd, dorsal auditory cortex; AUDp, primary auditory cortex; PH(d) (dorsal) posterior hypothalamus; GU, gustatory cortex; ENTDL, dorsolateral entorhinal cortex; PRh, perirhinal cortex; PIR, piriform cortex; LHA, lateral hypothalamic area; MOs, secondary motor cortex; VAL, ventral anterior-lateral complex; VPM, ventral posteromedial nucleus; VPL, ventral posterolateral nucleus; PO, posterior complex; CM, centromedian nucleus; VM, ventromedial nucleus; LA, lateral amygdala; CEA, central amygdala; MPO, medial preoptic area; SSs, secondary somatosensory cortex. The r in the first letter indicates right hemisphere and no small first letter indicates the left hemisphere.

In the correlation coefficient of selected features, all 10 features showed a significant reduction in correlation coefficients under anaesthesia compared to those in the awake state (Table 1). In accordance with the considerable effect size, interhemispheric connectivity in the anterior cingulate cortex and dorsomedial hypothalamus was extracted as a key feature. Additionally, a decrease in interhemispheric connectivity in the dorsolateral entorhinal cortex and lateral amygdala, which showed between medium (between 0.5 and 0.8) and large effect sizes (greater than 0.8), was observed. It should be noted that eight of ten features estimated from LASSO showed medium or large effect sizes, although some were not included in the top 20 ranking of Cohen’s d (Fig. 4B). The connectivity between the left gustatory area and VPL and between the left CA3 and right temporal association area showed small effect sizes, which were between 0.2 and 0.4.

### Differences in dynamic FC under awake, Iso and Iso + Med anesthetized conditions

Dynamic FC was then investigated in the awake, Iso, and Iso + Med anaesthetized conditions. K-means clustering identified four distinct connectivity states based on the windowed FC matrices (Fig. 5A). State 1 was characterized by strong positive and negative FC within the LCN. State 2 was characterized by widespread positive and negative FC. State 3 was characterized by overall weak FC. State 4 was characterized by dominant positive FC within the LCN, Hypo and AUD-VIS and negative FC between the LCN and Hypo. The dwell time window was between 9 and 100 in the awake state (Fig. 5B). The dwell time window of state 1 significantly increased under Iso + Med anaesthesia compared to that of the awake state, whereas the dwell time window under Iso anaesthesia increased from that of the awake state but not significantly (5.56 ± 3.13 for awake state, 26.34 ± 9.61 for Iso and 144.86 ± 44.63 for Iso + Med). In contrast, the dwell time window in state 4 under Iso and Iso + Med anaesthesia significantly decreased compared to that of the awake state (33.19 ± 10.55 for awake state, 4.59 ± 2.62 for Iso and no dwell time was found for Iso + Med). The frequency of occurrence in state 1 increased under Iso (0.27 ± 0.09) and Iso + Med anaesthesia (0.75 ± 0.10) compared to that of the awake state (0.03 ± 0.02), whereas the frequency of occurrence in state 4 was decreased under Iso (0.02 ± 0.02 for state 4) and Iso + Med (no occurrence was observed for state 4) anaesthesia compared to that of the awake state (0.30 ± 0.10) (Fig. 5C). Together with the dwell time results, this finding indicates that the anaesthetized state is characterized by the shift of dynamic FC from state 4 to state 1.

**Figure 5.**
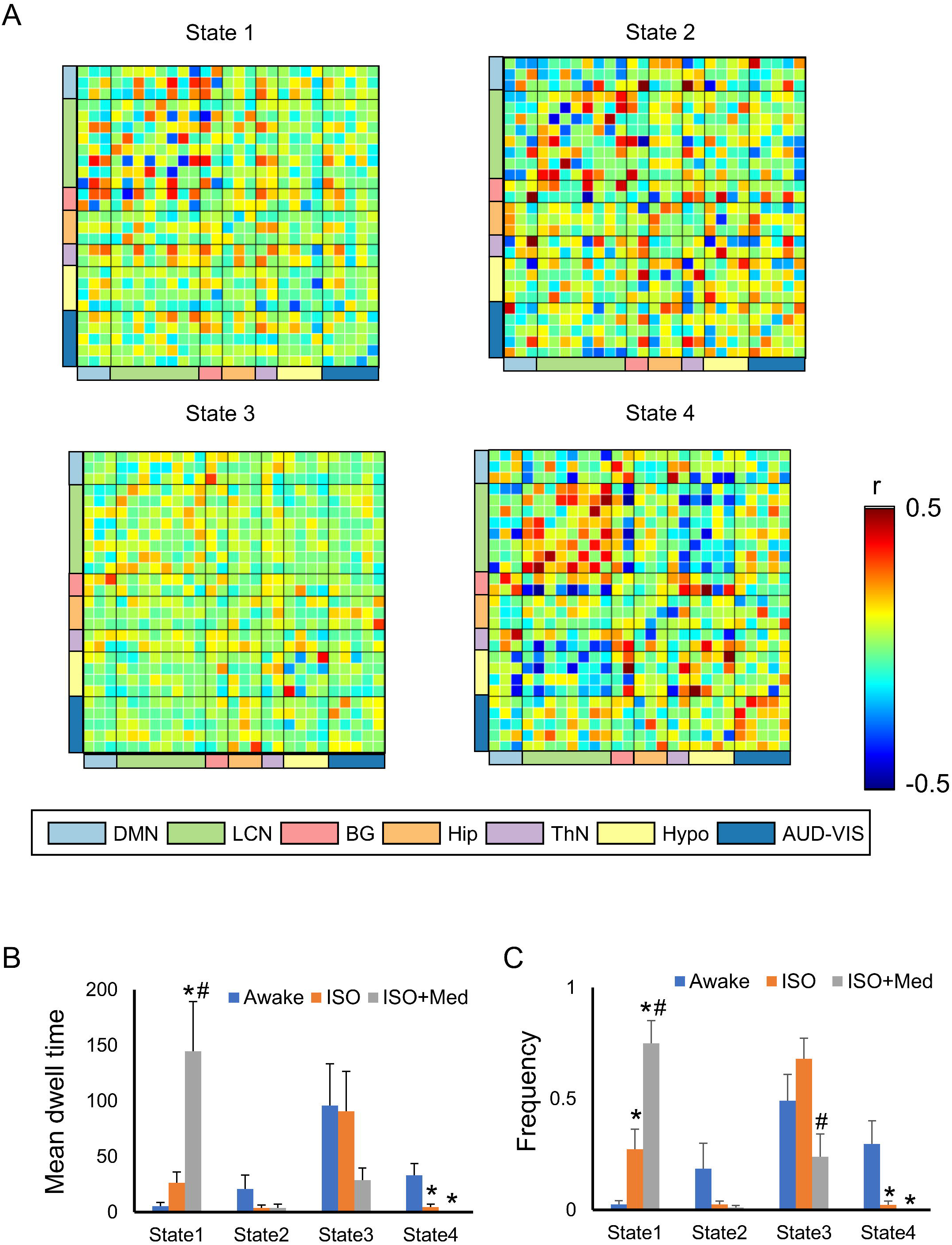
Dynamic functional connectivity in awake, Iso anaesthetized and Iso + Med anaesthetized mice. (A) Four dynamic FC states. (B and C) Averaged dwell time (B) and frequency of occurrence (C) in each state in the awake condition, under Iso anaesthesia and under Iso + Med anaesthesia. *p<0.05 compared with the awake condition in each state, Bonferroni-corrected. ^#^p<0.05 compared with the Iso condition in each state, Bonferroni-corrected.

### Alterations in fALFF under Iso and Iso + Med anaesthesia compared with the awake state

We compared fALFF between awake and anaesthetized states. The fALFF in the medial part of the thalamus, such as the CM, significantly decreased under Iso anaesthesia, while the fALFF in the hypothalamus increased under Iso anaesthesia (Fig. 6A). Iso + Med anaesthesia significantly decreased fALFF in several thalamic nuclei, including the CM and VPM/VPL (Fig. 6B). The fALFF in the cortex, including the anterior cingulate cortex (ACC), primary somatosensory cortex (SSp), auditory cortex (AUD), and entorhinal cortex (ENT), and the hypothalamus, including the medial preoptic area (MPO) and ventromedial nucleus (VMH), significantly increased under Iso + Med anaesthesia compared with the awake state (Fig. 6B). There was no significant difference in fALFF between Iso and Iso + Med anaesthesia except for a small spot of left SC (Supplementary Fig. 3A). We also compared fALFF between awake and anaesthetized (combining Iso and Iso + Med) states (Fig. 6C). fALFF in the thalamic nuclei, including the CM and VPM, was significantly decreased, and fALFF in the cortex, including the motor cortex, somatosensory cortex, auditory cortex and entorhinal cortex, and the hypothalamus was increased under anaesthetization. The test-retest reproducibility in fALFF in awake mice was assessed. There was no significant change in fALFF between the first and second scans except for the small area showing an increase in the cortex in the second scan (Supplementary Fig. 3B). Together with the static FC results (Supplementary Fig. 1), awake fMRI shows excellent reproducibility. (Figure 6)

**Figure 6.**
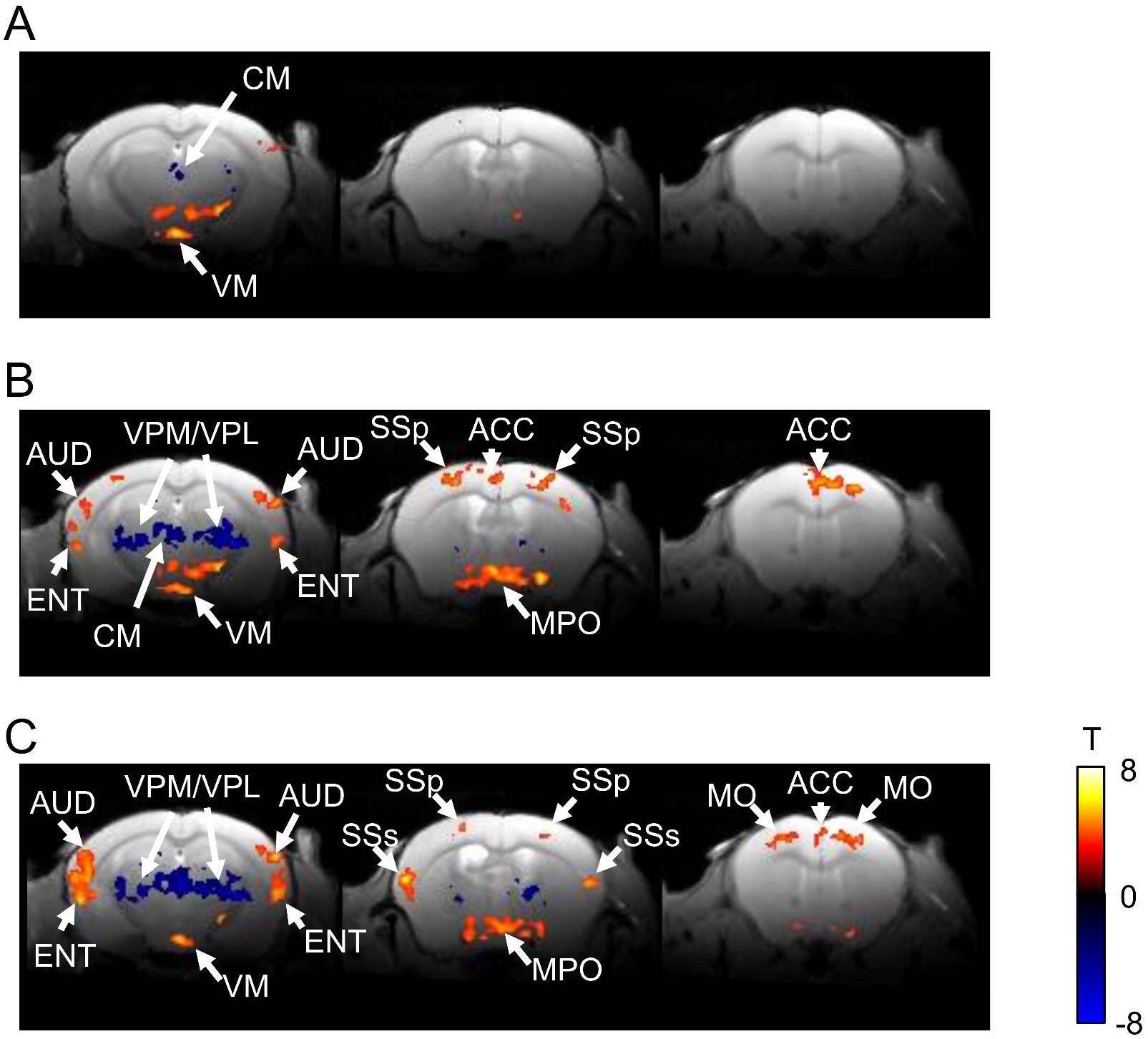
fALFF in awake, Iso-anaesthetized and Iso + Med-anaesthetized mice. (A) Significant increase in fALFF (hot colour) and decrease in fALFF (blue colour) in the awake group compared with Iso group. (B) Significant increase in fALFF (hot colour) and decrease in fALFF (blue colour) in the awake group compared with Iso + Med group. (C) Significant increase in fALFF (hot colour) and decrease in fALFF (blue colour) in the awake group compared with the anaesthetized group. P < 0.05, FDR-corrected. Colour bar T-value. ACC, anterior cingulate cortex; AUD, auditory cortex; ENT, entorhinal cortex; MPO, medial preoptic area; VM, ventromedial nucleus; MO, motor cortex; SSp, primary somatosensory cortex; SSs, secondary somatosensory cortex; VPL, ventral posterolateral nucleus; VPM, ventral posteromedial nucleus.

## Discussion

In this study, we investigated the alteration of static and dynamic functional connectivity in awake state and anaesthetized state in the same animals. Most previous studies compared the anaesthetized state and awake state in different animal groups, which were scanned on different days. We successfully conducted the awake-anaesthesia fMRI protocol to identify shift in the functional network from awake to anaesthesia. Remarkably, awake fMRI will reduce the number of animals, suggesting the contribution to the 3R for animal experiment. Furthermore, dynamic functional connectivity indicates that dynamic FC shifted from positive and negative connectivity across LCN, Hypo and AUD-VIS (state 4 in Fig. 5) to dominant connectivity in the LCN (state 1 in Fig. 5). Static FC demonstrates a reduction in interhemispheric connectivity as well as in the subcortical network. Together, reduction of the FC in the subcortex, including the thalamic nuclei, limbic system and hypothalamus, is a key feature for the anaesthetized state.

### Awake fMRI and anaesthetized fMRI

Previous studies used different mouse groups to compare the effect of anaesthetics on static FC ^9, 10, 11^. Because resting-state FC architecture has individual variability ^29^, direct comparison of anaesthesia in the same animal could directly demonstrate FC in awake and anaesthetized states without individual variation confounding the results. The similarity across the subjects showed robust static FC in the awake state, whereas the anaesthetized state showed lower similarity. It is opposite to our presumption that animals would exhibit uniform FC under anaesthesia, but it is explained by that residue of physiological noise could be more effective on weak FC under anaesthesia. The current study also showed a similar reduction in static FC between Iso and Iso + Med. This is in contrast to previous studies showing robust reduction of static FC under Iso anaesthesia rather than Iso + Med anaesthesia ^11, 12^. Generally, Iso suppresses neurovascular coupling ^32^ and astrocyte activity as well as neuronal activity in a dose-dependent manner ^33, 34^. Our previous study also demonstrated that the Iso dose-dependently influenced the relationship between spontaneous neuronal activity and BOLD signals ^35^. A static FC study revealed that Iso dose-dependently induces a widespread reduction in static FC ^36^. Paasonen *et al* used 1.3% Iso for the Iso group and 0.06 mg/kg/h MED and 0.5–0.6% ISO for the Iso + Med group ^11^. They reported lowed static FC under Iso anaesthesia because higher dosage of Iso was used. Here, we used 0.8% and 0.5% Iso, which was enough to observe spontaneous neuronal activity and BOLD fluctuation, for Iso and Iso + Med anaesthesia condition respectively. This low dose of Iso did not matter for static FC. Importantly, dynamic FC and fALFF showed distinct changes between Iso and Iso + Med anaesthesia. As described below, Med itself modulates neuronal activity through α2-adrenoreceptors, which indicates that dynamic FC and fALFF are more sensitive to the modulation of neuronal activity and temporal synchronization between neurons affected by Med.

### Static and dynamic functional connectivity

In this study, a comparison was made between awake and anaesthetized conditions using static FC and dynamic FC measures. We observed a widespread reduction in static FC, and interhemispheric connectivity was reduced both under Iso and Iso + Med anaesthesia, consistent with a previous study ^11, 37^. Subcortical regions, including the agranular insular cortex, thalamic nuclei and a part of the limbic system, are mostly included in the reduction of static FC. These results are consistent with previous studies in humans showing that static FC of the thalamus preferentially decreased under propofol and dexmedetomidine anaesthesia in humans ^4, 38, 39^. The anterior insular cortex is an informational hub in the global network related to consciousness ^40^. Previous rodent studies compared static FC between awake and anaesthetized states ^11, 37^,

but the ROI size was too large to investigate the subnucleus level. Our static FC study, therefore, provides the first evidence that investigated different brain networks between the awake state and anaesthetized state in detail. High-resolution fMRI showed that several nuclei of the thalamus were part of key connections in the anaesthetized state (Fig. 3). In addition to comparing static FC between the awake and anaesthetized groups, we used LASSO, which is a linear regression technique enforcing sparsity on its weights, to extract key connectivity by machine learning. Sparsity has an advantage that creates interpretable results by selecting a small number of important features. We found several key connections from those that showed large Cohen’s d values. This indicates the feasibility of LASSO to extract the essential FC for alteration of brain network from all connections that had moderate-large effect size. However, very few connections that showed small effect sizes were estimated as key features by LASSO. We should carefully assess the LASSO results to validate the key structure.

Dynamic FC offers an advantage in assessing dynamic functional network because connectivity is not stable during measurements. In addition to this advantage, dynamic FC is useful to investigate brain connectivity of individuals ^41^. We observed functional networks such as the DMN, LCN, BG, AUD-VIS, Hip, ThN and Hypo, which are commonly identified in rodent ICA studies ^12, 42^. The dynamic FC study demonstrated that the frequency of occurrence within the LCN (state 1) increased under anaesthesia, and instead, interaction within the LCN, Hypo and VIS-AUD (state 4) decreased. We inferred that a reduction in connectivity strength in the subcortical regions could be related to reduction of the dynamic interaction of the global functional network. Remarkably, Iso + Med group significantly reduced weak global interaction (state 3) rather than Iso group, indicating mixture of Iso and Med induced more significant reduction of global network than Iso only. These results indicate that the switching of the frequency of occarance between local and distant networks is the key mechanism underlying shifts between awake and anaesthetized states.

### fALFF increase and neuronal activity

The fALFF in the medial nuclei of the thalamus, including the centromedian nuclei, was significantly decreased under Iso and Iso + Med anaesthesia (Fig. 5). This corresponds to our previous study showing that the activity of the centromedian nuclei of the thalamus is related to the arousal state in rats ^43^. In a previous study, we clearly showed that the apparent diffusion coefficient, which was calculated from diffusion MRI signals, was related to neuronal activity. The ADC as well as the local field potential in the CM depends on the arousal level. Here, in addition to ADC, we showed that fALFF is another method to track the neuronal activity. Indeed, we showed that fALFF was correlated to neuronal activity in mice ^21, 30^. Importantly, because fALFF can be computed with the same data set for FC, it is possible to compare fALFF and FC directly in the same group. The simultaneous measurement of neuronal activity by means of fALFF and synchronization of neuronal oscillation between anatomically separated regions will provide insights into what happens in the resting state during anaesthesia. In addition to the reduction in fALFF in the medial part of the thalamus, Iso + Med reduced fALFF in the lateral part of the hypothalamus. Iso suppresses neuronal activity in the CM ^44^, and medetomidine suppresses neuronal activity in the thalamus through the α2-adrenoreceptor, which is abundantly distributed in the thalamus ^45, 46^. The mixed administration of Iso and 12 Med therefore induces a stronger and broader reduction in neuronal activity in the thalamus than Iso alone. An [18F] FDG□PET study revealed an increase in glucose metabolism in the hypothalamus under Iso anaesthesia, supporting our results that fALFF in the hypothalamus, including the POA, increased both under Iso and under Iso + Med anaesthesia ^47^. Dexmedetomidine, an analogue of medetomidine, increased the delta frequency power of local field potentials in the cortex ^48^. This finding can explain why fALFF in the cerebral cortex partially increased under Iso + Med anaesthesia.

### Perspectives and Conclusions

In the present study, we demonstrated alterations in static FC and dynamic FC under two types of anaesthesia compared with the awake state. The typical structure of static FC was conserved under anaesthesia, although the strength of static FC decreased compared to that observed in the awake state. The dynamic property of the functional network shifted to regional connectivity within the cortex under an anaesthetized state. These results indicate that connectivity in the subcortex could be a marker of the anaesthetized state. These results provide insight into the alteration/non-alteration of neuronal networks under unconscious conditions.

## Acknowledgements

We thank to Mr. Boucif Djemai for the support of MRI experiment.

## Funding sources

This research in the laboratory of TT was supported by *Fondation de France* (grant number: 00086313) and *Grant-in-Aid for Research Activity Start-up* (grant number: 20K22698).

## CRediT authorship contribution statement

**Tomokazu Tsurugizawa:** Conceptualization, Methodology, Investigation, Visualization, Writing - review & editing. **Daisuke Yoshimaru:** Methodology, Visualization, Writing - review & editing.

## Figure legends

**Supplementary Figure 1. Test-retest reproducibility and comparison of functional connectivity between anaesthetized states**.

(A) Averaged connectivity matrices in the first scan and second scan of awake groups. (B) Statistical significance of test-retest data. The lower left shows the T-values, and the upper right shows the significant differences (p < 0.05, NBS-corrected). (C) Comparison between Iso and Med groups. The lower left shows the T-values, and the upper right shows the significant differences (p < 0.05, NBS-corrected).

**Supplementary Figure 2. Common ICA components in awake, Iso-anaesthetized and Iso + Med-anaesthetized states by group ICA based on 40 components**.

Common ICA components, default mode network (DMN), lateral cortical network (LCN), subcortical network (BG), hippocampus (Hip), thalamus (ThN), Hypothalamus (Hypo), auditory-visual cortical network (AUD-VIS) are overlaid on the representative mouse image.

**Supplementary >Figure 3. Comparison of fALFF between the anaesthetized state and fALFF of test-retest reproducibility of awake fMRI**.

(A) Significant increase in ALFF (hot colour) and decrease in ALFF (blue colour) in the Iso group compared with the Iso + Med group. (B) There was no significant increase in ALFF (hot colour) or decrease in ALFF (blue colour) between the first scan and second scan in the awake group. P < 0.05, FDR-corrected. Colour bar T-value.

## Abbreviations

ACC: anterior cingulate cortex
AId: dorsal agulanular insular cortex
AIp: posterior agulanular insular cortex
AUDd: dorsal auditory cortex
AUDp: primary auditory cortex
AUD-VIS: auditory-visual cortical network
BOLD: blood oxygenation level dependent
CEA: central amygdala
CM: centromedian nucleus
DMN: default mode network
DMH: dorsomedial hypothalamus
ECT: ectorhinal area
ENTDL: dorsolateral entorhinal cortex
FC: functional connectivity
FD: Framewise displacement
FDR: false discovery rate
GU: gustatory cortex
Hypo: hypothalamic network
ICA: independent component analysis
Iso: isoflurane
LA: lateral amygdala
LASSO: least absolute shrinkage and selection operator
LCN: lateral cortical network
LHA: lateral hypothalamus
LN: limbic network
LSr: lateral septal nucleus
Med: medetomidine
MO(s): (secondary) motor cortex
MPO: medial preoptic area
MSE: minimum mean squared error
NBS: network based statistic
PAG: periaqueductal tract
PH: posterior hypothalamus
PHd: dorsolateral posterior hypothalamus
PIR: piriform cortex
PO: posterior complex
POA: preoptic area
SSs: secondary somatosensory cortex
Tea: temporal association area
ThN: thalamus
VAL: ventral anterior-lateral complex
VM: ventral medial nucleus
VPL: ventral posterolateral nucleus
VPM: ventral posteromedial nucleus

## Notes

### Competing Interest Statement

The authors have declared no competing interest.

