## Supplementary material for "Impact of anaesthesia on static and dynamic functional connectivity in mice": Table1

**Table 1 Discriminative features for each pair of connection from the feature-selection step of LASSO**

| Pair of connections | | P values | Effect size |
| --- | --- | --- | --- |
| Left Gustatory area | Left Ventral posterolateral nucleus of the thalamus | 0.0002 | 0.29 |
| Left anterior cingulate cortex | Right anterior cingulate cortex | < 0.0001 | 1.44 |
| Left ectorhinal cortex | Left entorhinal cortex | < 0.0001 | 1.02 |
| Left CA3 | Right temporal association areas | 0.0016 | 0.42 |
| Left entorhinal cortex | Right entorhinal cortex | 0.0016 | 0.79 |
| Left lateral amygdala nucleus | Right lateral amygdala nucleus | < 0.0001 | 0.62 |
| Left ventral anterior-lateral complex of the thalamus | Left ventral medial nucleus of the thalamus | < 0.0001 | 0.70 |
| Left ventral anterior-lateral complex of the thalamus | Right ventral posteromedial nucleus of the thalamus | < 0.0001 | 0.75 |
| Left paraventricular hypothalamic nucleus | Right paraventricular hypothalamic nucleus | 0.0002 | 1.20 |
| Left dorsomedial nucleus of the hypothalamus | Right dorsomedial nucleus of the hypothalamus | < 0.0001 | 1.29 |
